# Improving the consistency of functional genomics screens using molecular features - a multi-omics, pan-cancer study

**DOI:** 10.1101/322883

**Authors:** Wenyu Wang, Alina Malyutina, Alberto Pessia, Caroline A. Heckman, Jing Tang

## Abstract

Probing the genetic dependencies of cancer cells helps understand the tumor biology and identify potential drug targets. RNAi-based shRNA and CRISPR/Cas9-based sgRNA have been commonly utilized in functional genetic screens to identify essential genes affecting growth rates in cancer cell lines. However, questions remain whether the gene essentiality profiles determined using these two technologies are comparable. In the present study, we collected 42 cell lines representing a variety of 10 tissue types, which had been screened both by shRNA and CRISPR techniques. We observed poor consistency of the essentiality scores between the two screens for the majority of the cell lines. The consistency did not improve after correcting the off-target effects in the shRNA screening, suggesting a minimal impact of off-target effects. We considered a linear regression model where the shRNA essentiality score is the predictor and the CRISPR essentiality score is the response variable. We showed that by including molecular features such as mutation, gene expression and copy number variation as covariates, the predictability of the regression model greatly improved, suggesting that molecular features may provide critical information in explaining the discrepancy between the shRNA and CRISPR-based essentiality scores. We provided a Combined Essentiality Score (CES) based on the model prediction and showed that the CES greatly improved the consensus of common essential genes. Furthermore, the CES also identified novel essential genes that are specific to individual cell types. Taken together, we provided a systematic approach to define a more accurate gene essentiality profile by integrating functional screen data and molecular profiles.

## Introduction

Interrogating the genetic dependencies of cancer cells is a fundamental step in target-based drug discovery (Mullenders and Bernards, 2009). Loss-of-function genetic screens have emerged as powerful tools to introduce gene knockdown in vitro, providing novel insights into the genes that are essential for cell survival and proliferation, which are of pivotal values for target identification and validation (Fellmann *et al.*, 2017). To carry out a systematic exploration of cancer dependency profiles at the genome scale, these functional genetic screens rely on a pooled library containing tens of thousands of synthetized short sequence constructs, which are designed to target different genes. Using an optimal delivery system, the pooled library as a whole can be efficiently introduced into a cell culture, resulting in a mixture of cell subpopulations, each of which carries one sequence construct that triggers the depletion of a particular gene. During the culture period, the cell subpopulations depleted of essential genes will lose fitness, resulting in a diminished abundance of their effector sequence constructs (Mullenders and Bernards, 2009). The change in relative abundance of the sequence constructs can thus represent the degree of essentiality.

Over the last decade, shRNA (short hairpin RNA), together with the more recently developed CRISPR-Cas9-based sgRNA (single guide RNA) have been adopted as two major technologies to carry out pooled loss-of-function genetic screens. The experimental procedures for shRNA-based and CRISPR-based pooled screens are quite similar, while the major differences being the synthetic sequence constructs that are delivered into the cells to activate distinct loss-of-function machineries. shRNA is directed via the RNAi (RNA interference) pathway to bind to its target mRNA in the cytoplasm, leading to the degradation of the target mRNA without changing the genome of the cells (i.e. a transient knockdown effect). sgRNA, on the other hand, utilizes CRISPR pathway to direct the Cas9 protein to cut genomic DNAs in the nucleus, triggering the non-homologous end joining (NHEJ) pathway to introduce most often permanent loss-of-function mutations, resulting in a complete knockout of the target genes (Doench, 2018; Fellmann *et al.*, 2017; Morgens *et al.*, 2016). Despite the relative simplicity to set up the experiments, the efficiency and specificity of the sequence constructs have been limiting factors for a reliable detection of cancer essential genes. For example, evidence has been found that both shRNAs and sgRNAs may affect unwanted off-target genes with partial sequence complementarity, introducing noises that mask the actual growth phenotype of the intended gene depletions (Boettcher and McManus, 2015; Schaefer *et al.*, 2017). Differences in gene-depletion efficiency also contribute to the phenotypic variabilities for shRNA screens (Barrangou *et al.*, 2015) as well as CRISPR screens (Doench *et al.*, 2016). There have been considerable efforts to improve the design of the sequence constructs to achieve maximal efficiency and minimal off-target effects (Boettcher and McManus, 2015; Munoz *et al.*, 2016). Computational methods have also been proposed to filter out the off-target effects to improve the screen quality (Doench *et al.*, 2016). Furthermore, statistical methods have been developed to control false positives by evaluating multiple sequence constructs targeting the same genes (Barbie *et al.*, 2009; Li *et al.*, 2014).

Given the increasing maturity of both shRNA and CRISPR technologies to detect cancer essential genes, a logical next question would be: whether the essentiality scores from both screens are consistent for the same gene. A side-by-side comparison of the gene essentiality profiles at the genome scale should provide important insights about the advantages and limitations of the two screens. Two recent studies carried out shRNA and CRISPR screens in parallel for several human cancer cell lines (Evers *et al.*, 2016; Morgens *et al.*, 2016), with different conclusions being made in terms of their relative performances for detecting truly essential genes. Evers et al. reported a superior accuracy using CRISPR compared to shRNA, while Morgens et al. observed similar performances. Interestingly, Morgens et al. found that a big proportion of essential genes identified by one screen were not found in the other screen, suggesting complex confounding factors that are inherently distinct between these two technologies. However, their conclusions were made based on very limited data, as Evers et al. investigated the prediction accuracy for a small gene set including 46 essential and 47 non-essential genes using only two cancer cell lines (RT-112 and UM-UC-3), while Morgens et al. analyzed a much bigger gene set including 217 essential and 947 non-essential genes, but the comparison was done using only one cell line (chronic myelogenous leukemia cell line K562).

In this study, we carried out a more systematic comparison between shRNA and CRISPR data across a larger collection of cancer cell lines that were available from existing publications and databases. We found that the shRNA and CRISPR-based gene essentiality profiles showed limited consistency irrespective of the cancer types or the status of essentiality, even after adjusting the off-target effects using computational methods. On the other hand, we found that molecular features of genes can explain the discrepancy of the gene essentiality between the two screening technologies. By constructing a regression model with the molecular features as covariates, the accuracy of predicting gene essentiality can be greatly improved, suggesting that molecular information should be considered when evaluating gene essentiality scores. We developed a computational approach called Combined Essentiality Scoring (CES) to predict gene essentiality by integrating both functional and molecular features. We showed that CES can be used for predicting not only commonly essential genes across various cell lines, but also therapeutic targets that are specific for a certain cancer type. The CES scoring thus may serve as a novel informatics tool to facilitate the discovery of cancer gene dependency in personalized medicine.

## Materials and Methods

### Integrating functional screening and molecular profiling data

We aimed at a genome-wide comparison between shRNA- and CRISPR-based essentiality scores on multiple cancer cell lines. A total of 42 cancer cell lines with both shRNA and CRISPR screening data were collected from the Achilles database (v3.38 and v2.20) (Aguirre *et al.*, 2016; Tsherniak *et al.*, 2017) and three recent studies (Hart *et al.*, 2015; Steinhart *et al.*, 2017; Wang *et al.*, 2017) (**Table S1**). Gene-level shRNA-based essentiality scores were obtained by averaging over the sequence-level essentiality scores. The CRISPR-based gene essentiality scores were summarized using averaged (Wang *et al.*, 2017), second-best (Aguirre *et al.*, 2016) or Bayesian modeling averaging strategy (Hart and Moffat, 2016). The interpretation of essentiality scores was such that the lower the score the more important the gene is for cancer growth. We utilized the DEMETER approach to correct for the off-target effects in the shRNA screens (Tsherniak *et al.*, 2017). As the essentiality scores represent the fold change of the sequence constructs, we utilized Pearson correlation as the main metric, where a positive correlation was considered good consistency and a zero or negative correlation indicated poor consistency. We also used mean square error (MSE) and adjusted coefficient of determination (R^2^) to evaluate the prediction performance of different linear regression models.

Molecular profiles of the genes for these 42 cell lines, including mutation, gene expression and copy number variation were downloaded from the Cancer Cell Line Encyclopedia (CCLE) database (Barretina, et al., 2012). For the gene expression profiles, both the RNA-Seq-based RPKM counts (denoted as Exp.seq) and Affymetrix-based microarray intensities (denoted as Exp.array) were utilized. The resulting data matrix thus contains the two versions of essentiality scores and molecular profiles for each gene for a given cell line, wherever available (**Figure 1**). All the features were mean normalized as z-scores for further analyses, resulting in 386, 186 data points involving 14,138 genes. A detailed list of the data sources can be found in **Table S1**. For evaluating the prediction accuracy of the different prediction methods, a set of 3,804 housekeeping genes was collected from a previous study as the ground truth for the common essential genes across all cell types (Eisenberg and Levanon, 2013) (**Table S2**). The pathway enrichment analyses for consistent versus inconsistent gene sets were done using the GSEAPreranked method (Subramanian, et al., 2005) and the Enrichr web server (Kuleshov *et al.*, 2016). Statistical significance was derived using specific hypothesis testing mentioned in the **Result** section.

**Fig. 1.**
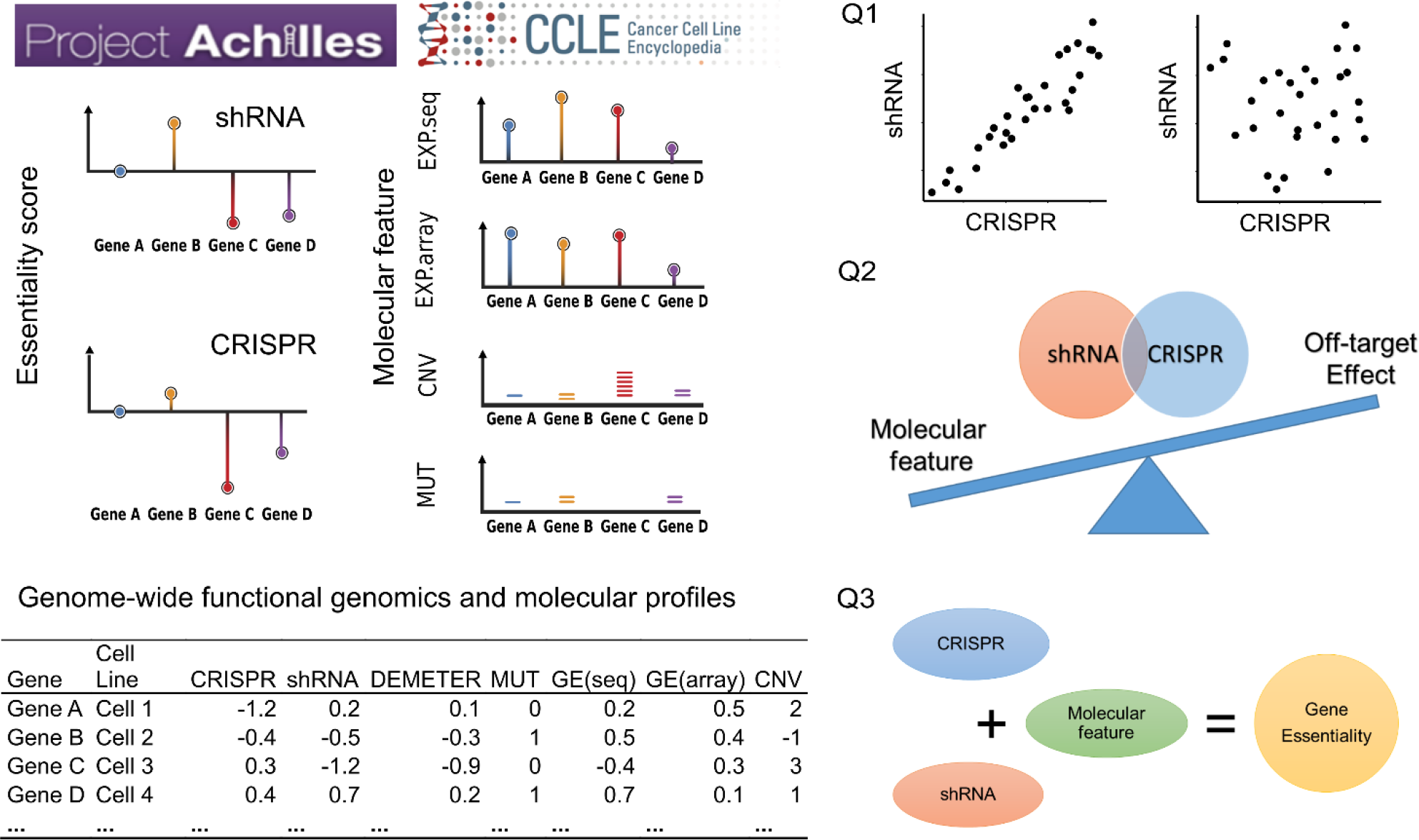
The data integration pipeline to evaluate the consistency of gene essentiality profiles from shRNA- and CRISPR-based screens. Functional screen data as well as molecular profiling data for each cell line at the genome scale were integrated from the public databases including the Achilles and CCLE data portals. For a gene at a cell line, a vector of 7 features was determined including CRISPR-based essentiality score, shRNA-based essentiality score, DEMETER corrected shRNA essentiality score, mutation status, RNA-seq-based gene expression, microarray-based gene expression and copy number variation. Three questions were investigated here: Q1) whether the shRNA and CRISPR-based gene essentiality scores are consistent; Q2) whether the molecular profiles including mutation, gene expression and copy number variation data can explain the differences between shRNA and CRISPR-based gene essentiality scores better than off-target effects; and Q3) whether there is a predictive model that can help improve the consensus of essential genes despite the technical and experimental biases.

## Results

### Poor consistency between shRNA and CRISPR-based gene essentiality scores

To evaluate how the choice of technology affects the gene essentiality scoring, we first checked the consistency of the gene essentiality scores determined by shRNA and CRISPR screens. We found generally poor consistencies across all 42 cell lines, where 25 showed positive but moderate correlations while 17 cell lines had zero or even negative correlations (average correlation = 0.04) (**Figure 2A**). The HT29 cell line (colon cancer) showed the maximal consistency with a correlation of 0.24. In contrast, we observed much poorer consistency for cell lines such as HPAFII (pancreatic cancer), DLD1 and HCT116 (colon cancers), with correlations decreasing to −0.17 and lower. In general, we did not find the enrichment of certain cancer types in the consistency ranking of the cell lines. However, 9 out of 10 leukemia cell lines have shown positive correlations, suggesting that leukemia cells tend to be more robust to the choice of technologies compared to the other cancer types. On the other hand, the only leukemia cell line that has a zero or negative correlation (−0.01) was K562, for which the poor consistency between shRNA and CRISPR screens was also confirmed in a side-by-side comparison by Morgens et al. (Morgens *et al.*, 2016). We also tested the accuracy of using the shRNA score to predict the CRISPR score. The mean squared errors (MSE) were not significantly different from that for a random prediction, confirming the poor consistency (p-value = 0.08, Wilcoxon test, **Figure 2B**).

**Fig. 2.**
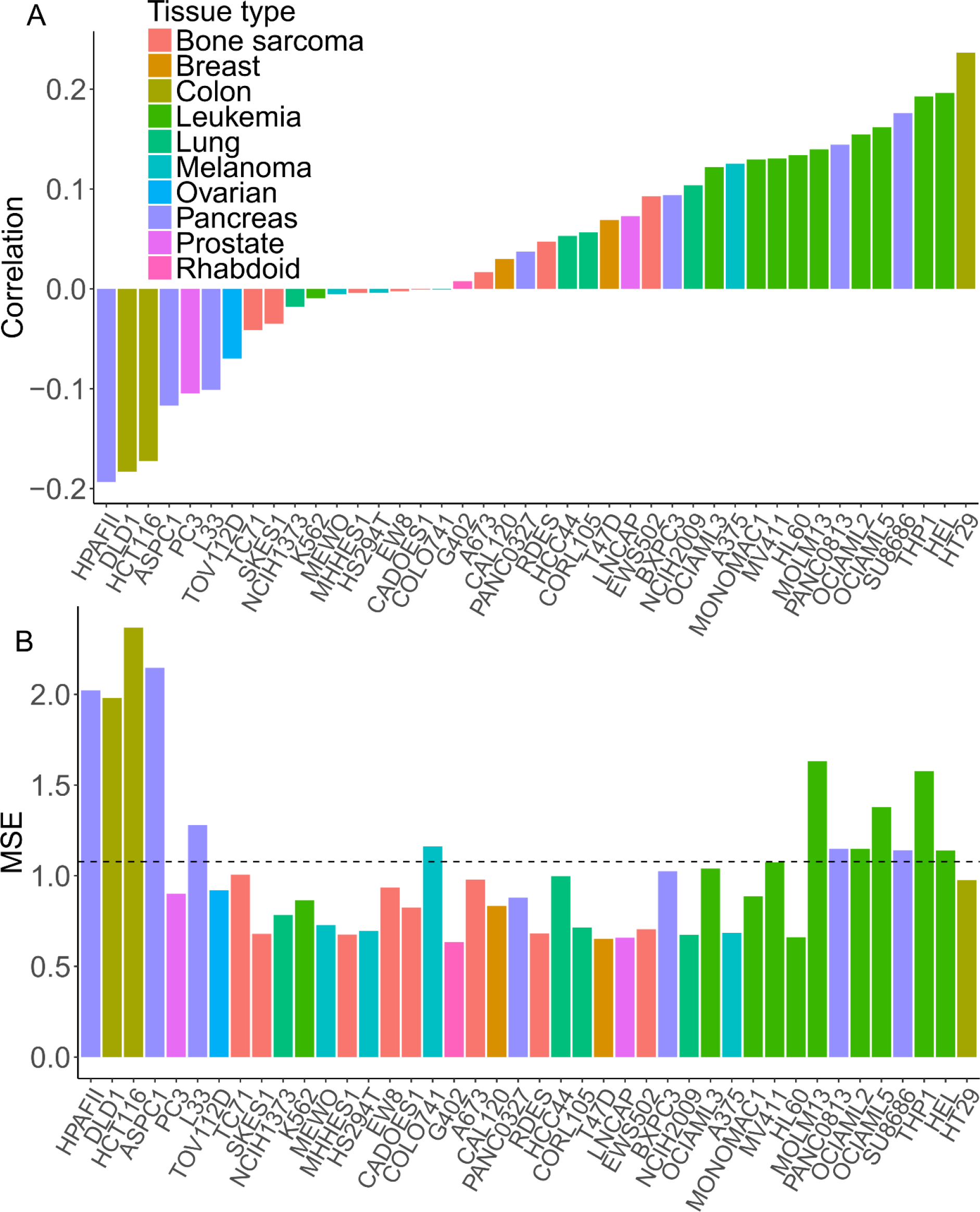
Poor consistency of the gene essentiality scores across a total of 42 cancer cell lines. (A) Pearson correlation between the shRNA and CRISPR scores (B) Mean squared error (MSE) using the shRNA score to predict the CRISPR score. The cancer types were shown in different colors. The average MSE for a random prediction was shown as the dashed line (MSE = 1.08).

Note that the gene-level essentiality scores were averaged over multiple sequence constructs for both the shRNA and CRISPR screens, therefore, the poor consistency could not be explained by the biases of certain sequence constructs. On the other hand, we found that both shRNA and CRISPR showed stronger essentiality scores for the set of housekeeping genes (n = 3,804) than non-housekeeping genes (Wilcoxon test, p-value < 2×10^−16^), suggesting the overall validity of the genome-wide functional screens. However, the consistency between the two screens based on the housekeeping genes seems to improve only for the 25 cell lines that show overall positive correlations (average correlation of 0.14 for the housekeeping genes versus 0.04 for the others). In contrast, for cell lines with overall negative correlations (n = 15), the consistency within the housekeeping genes became even lower (average correlation of −0.08 versus −0.02). These results suggested that the housekeeping genes did not necessarily show higher consistency between the shRNA and CRISPR screens than the other genes.

Next, we sought to evaluate the impact of off-target effects on the discrepancy between shRNA and CRISPR essentiality scores. Off-target effects due to seed-sequence complementarity have been known to confound shRNA screens. On the other hand, CRISPR is considered as a more specific genome-editing technique with much fewer off-targets (Zhang *et al.*, 2015). Therefore, we hypothesized that by adjusting for the off-target effects in shRNA screen results the consistency may be improved between shRNA- and CRISPR-based essentiality scores. We applied the recently developed DEMETER method to detect off-target effects by modeling the likelihood of seed-sequence complementarity of shRNAs (Tsherniak *et al.*, 2017). Surprisingly, the consistency across the cell lines did not improve significantly after the DEMETER correction (average increase of correlations = 0.02, p-value = 0.1, Wilcoxon signed rank test, **Figure 3A**). We observed a moderate increase of correlations for 30 cell lines, but not for the other 12 cell lines. The biggest improvement was observed for the PC3 cell line (prostate cancer), where the correlation was changed from −0.11 to 0.07 (an increase of 0.18). For the HT29 cell line (colon cancer) for which the original correlation was 0.24, the DEMETER correction managed to improve it to 0.37 (an increase of 0.13). In contrast, for the HL60 (leukemia) cell line, DEMETER correction decreased the consistency from a correlation of 0.13 to −0.09 (a decrease of −0.22). Interestingly, the decrease of consistency was observed for 9 leukemia cell lines, while the only leukemia cell line that showed improved consistency was K562 (**Figure 3A**). The correlations between the DEMETER score and the shRNA score were similar for leukemia cell lines and the other cell lines (average correlations = 0.54 versus 0.52, p-value = 0.29, Wilcoxon test, **Figure 3B**), suggesting that the DEMETER correction was done similarly across all the cell lines. Additional investigations were thus required to understand why the DEMETER correction decreased the shRNA-CRISPR consistency particularly in leukemia cell lines. After removing all the 10 leukemia cell lines, the improvement of the screen consistency became significant, while the effect size was rather marginal (average increase of correlation = 0.07, p-value = 1.7×10^−7^, Wilcoxon signed rank test). Taken together, the majority of the differences between the shRNA and CRISPR essentiality profiles could not be simply attributed to the off-target effects in shRNA screens.

**Fig. 3.**
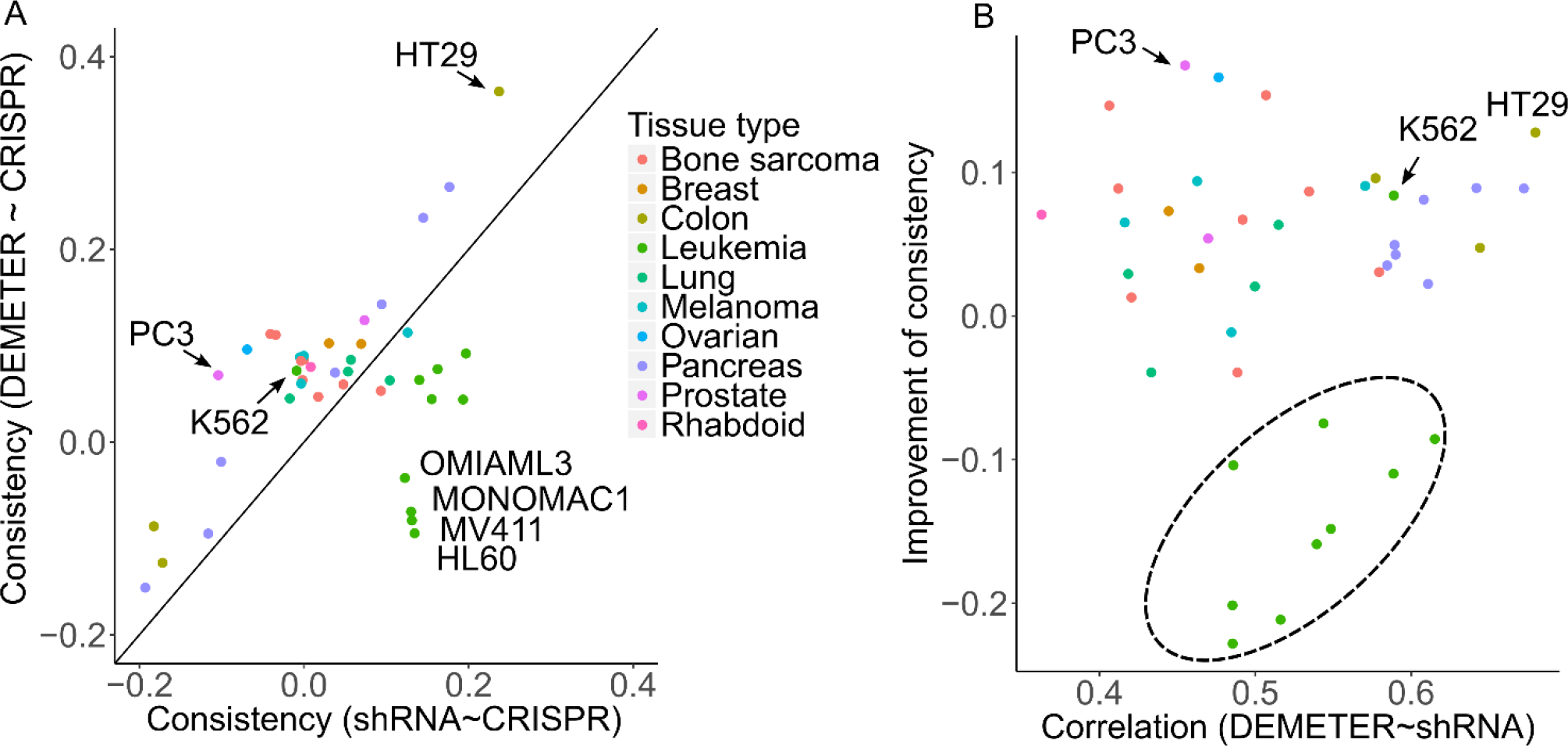
The consistency between shRNA and CRISPR screens after correcting the off-target effects. (A) The consistency before and after the DEMETER correction for each cell line. Example cell lines with improved as well as decreased consistency were labeled. (B) The majority of leukemia cell lines (highlighted in the circled area) showed decreased consistency even though the level of DEMETER correction was similar to the other cell lines.

Recent studies have shown that cells may respond to shRNA and CRISPR-Cas9 perturbations by activating distinct compensation mechanisms (El-Brolosy and Stainier, 2017; Peretz *et al.*, 2018). As a result, these defense mechanisms may interact with the pathways of the target genes, resulting in a confounding factor that affects the true essentiality profiles. Therefore, the consistency of shRNA and CRISPR screens at the pathway level may provide insights on which pathways tend to be affected by the compensation mechanisms. For example, if a pathway tends to be activated by the compensation mechanisms involved in shRNA but not in CRISPR (or vice versa), then the consistency of the genes within that pathway is expected low. Likewise, the pathways that include high consistency genes will be less biased and thus become more robust to different screen techniques. For example, we found that n = 10 genes involved in mRNA 3’-UTR binding (GO: 0003730) showed high consistency (average correlation = 0.41, **Figure 4A**), as compared to n = 64 genes in GPCR taste receptor activity (GO: 0090681) that showed little consistency (average correlation = −0.15, **Figure 4B**). For the n = 546 genes that showed average correlation higher than 0.3, we found n = 273 GO terms that were enriched. The top ones included DNA damage response, signal transduction by p53 class mediator (GO: 0030330) and thiol-dependent ubiquitin-specific protease activity (GO: 0004843) (adjusted p-value = 0.0005 and 0.002, respectively) (**Figure 4C**). In contrast, the GO terms specifically activated in low consistency genes (defined as those with average correlation < 0) were more versatile, ranging from GPCR-related pathways (e.g. GPCR taste receptor activity (GO: 0090681, adjusted p-value = 6.57×10^−7^) to calcium ion sensor activity (GO: 0061891, adjusted p-value = 1.28×10^−6^) (**Figure 4D**). These results suggested that the poor consistency between the shRNA and CRISPR screens are mainly driven by those genes with specific biological processes that may be activated unexpectedly by shRNA or CRISPR-Cas9 perturbations, which are however independent of the true essentiality of the intended target genes. These selective biological processes that are enriched in low consistency genes are worth further exploration with in-depth biological validations (**Table S3**).

**Fig. 4.**
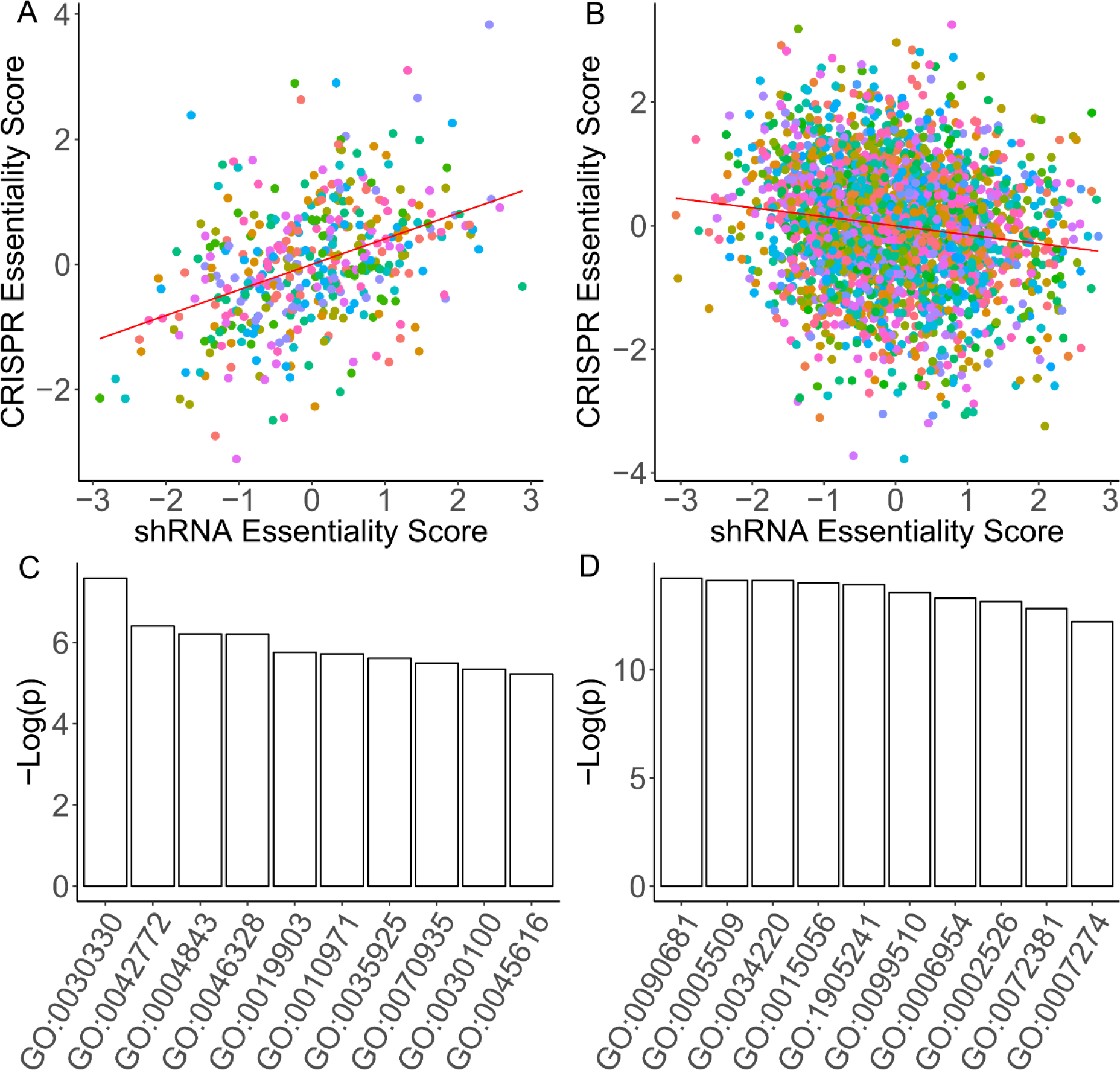
Genes that are involved in different biological processes showed opposite consistency patterns. (A) The consistency between shRNA and CRISPR screens for genes involved in mRNA 3’-UTR binding (GO: 0003730) as compared to those involved in (B) GPCR taste receptor activity (GO: 0090681). Colors of the data points correspond to the cell line identity. (C) The top 10 biological processes that are selectively enriched in high consistency genes as compared to (D) low consistency genes.

### Incorporating molecular features improved the consistency between shRNA- and CRISPR-based gene essentiality profiles

As the poor consistency between the shRNA and CRISPR gene essentiality profiles could not be explained by the off-target effects, and that the biological processes of genes seem to play a role, our next question became: can the molecular features of the genes, which are inherently determining their biological functions, bring additional information in explaining the missing consistency? This question could be answered quantitatively by formulating a regression problem to consider the CRISPR gene essentiality score as the response variable, to be estimated using shRNA gene essentiality score and molecular features including gene expression, mutation and copy number variation. The hypothesis thus became that if the molecular features contain useful information in explaining the differences between the shRNA and CRISPR screens, then the prediction accuracy of CRISPR score should be improved, compared to the use of shRNA-based scores alone. More specifically, we sought a comparison of the two regression models below.

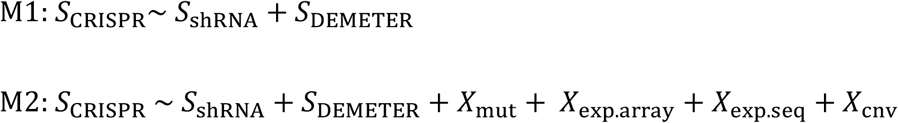

We considered separately the raw shRNA and the DEMETER-corrected essentiality scores, as they showed context-dependent performance when comparing to the CRISPR essentiality scores (**Figure 3**). Note that M1 was implicitly tested by the correlation analyses described in **section 3.1**, where it predicted the CRISPR-based score with limited accuracy. Here we applied a linear regression model M2 by adding molecular features including mutation, copy number variation, array-based and sequence-based gene expression values.

For each cell line, we adopted a 10-fold cross-validation to train the model using 70% of the data and test the model predictions on the remaining 30% of the data. The process was repeated 20 times to allow a random sampling of training and testing data. As shown in **Figure 5A**, M2 outperformed M1 significantly across all the cell lines, with the average correlation nearly doubled from 0.15 to 0.28 (p-value = 2.5×10^−8^, Wilcoxon signed rank test). The adjusted coefficient of determination was also increased three folds from 0.03 to 0.09 (p-value = 2.5×10^−8^, Wilcoxon signed rank test). When we randomly shuffled the molecular profiles of the genes and reran the M2 model, the prediction accuracy decreased to the same level as that of M1 (denoted as M2.perm in **Figure 5A**), suggesting that the actual molecular features are critical for contributing the improved performance of M2. However, when investigating the M2 model at the individual cell lines, we found that the performance varied (**Figure 5B-C**). For example, the molecular features helped achieve a correlation of 0.45 from 0.23 for a leukemia cell line MV411 (an increase of correlation = 0.22), while for a pancreatic cancer cell line (PAC0327) the improvement was rather marginal (an increase of correlation from 0.07 to 0.10). Similar trends were also observed for R2 and MSE (**Figure S1**). Therefore, the molecular features seemed to play a cell-dependent role in explaining the difference between the CRISPR and shRNA essentiality scores.

**Fig. 5.**
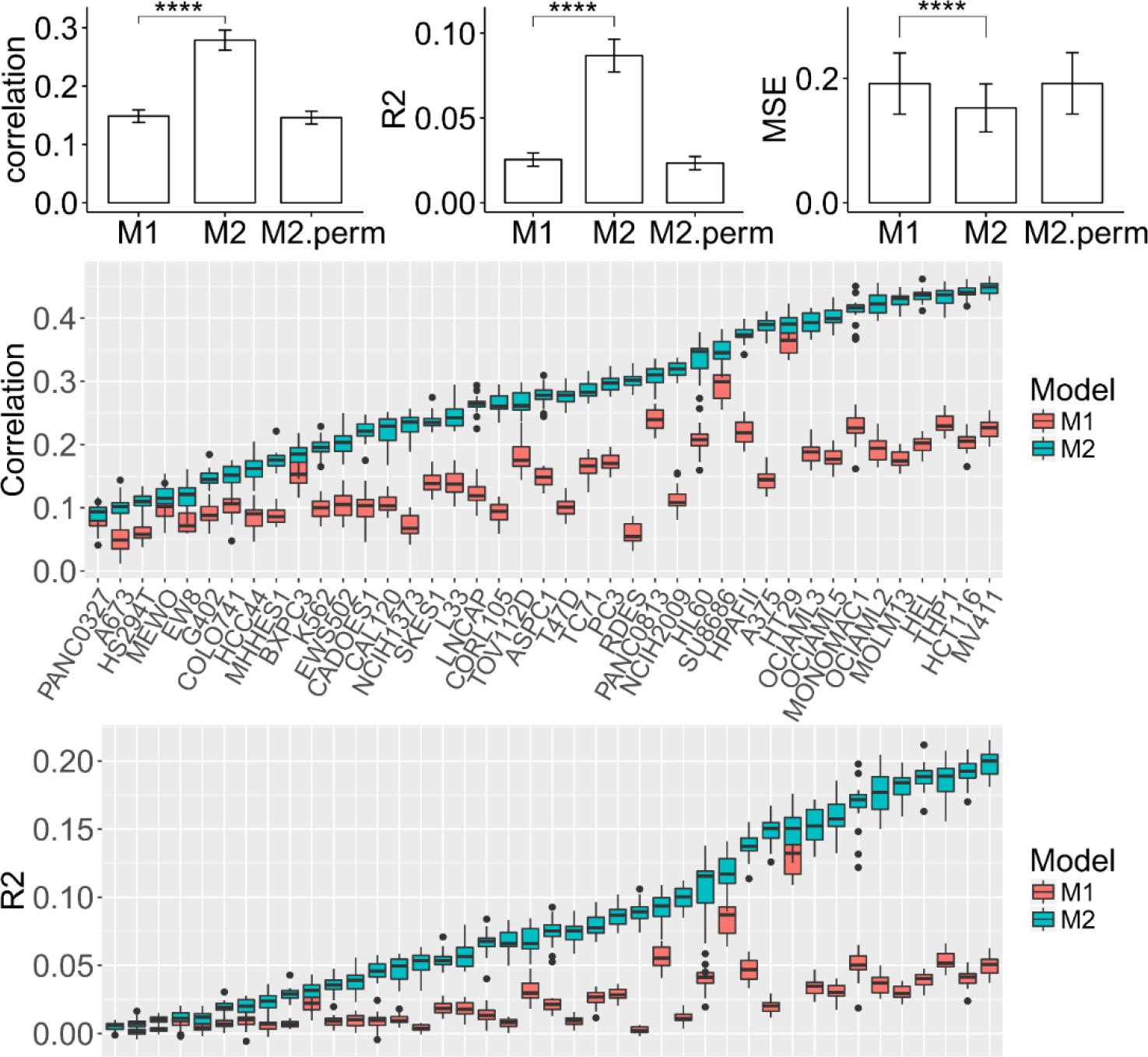
Model performance in terms of correlation and adjusted coefficient of determination and mean squared error for (A) all and (B-C) individual cell lines. ****: p < 0.0001, Wilcoxon signed rank test. M1: the linear regression model considering shRNA and DEMETER scores as explanatory variables; M2: the multiple linear regression model by including molecular features as additional covariates; M2.perm: the linear regression model where the molecular features were permutated.

Furthermore, we did the model comparison at the gene level using the same strategy and found that the same conclusion can be made, that for most of the genes, M2 outperformed M1 as the molecular features improved significantly the prediction accuracy and thus helped explain the difference between shRNA and CRISPR essentiality scores (**Figure S2**). The M2 model that was fitted at the gene level was easy to interpret, as the regression coefficient of a given molecular feature indicates its relative importance to explain the difference between the shRNA and CRISPR essentiality scores (**Table S4**). For example, for the *WRN* gene, which is known to maintain the DNA structure and stability, poor consistency was found between shRNA and CRISPR screens (correlation = −0.41), while M2 can predict the CRISPR scores with a much improved correlation of 0.72 (**Figure 6A**). The most significant regression coefficient was for mutation (coefficient = 0.6, p-value = 4.3×10^−5^), suggesting that the mutation status of *WRN* has been an important predictor for its CRISPR-based essentiality score (**Figure 6B**). Indeed, we found a positive correlation between the *WRN* mutation and CRISPR-based essentiality score (correlation = 0.67), suggesting that wildtype *WRN* was more essential than mutant *WRN*. However, the opposite conclusion could be made from the shRNA-screen, as the correlation between *WRN* mutation status and shRNA-based score turned negative (correlation = −0.40). For example, the CRISPR screen showed that inhibiting *WRN* in the HCT116 cell line (colon cancer) stimulated the cell growth, resulting in a positive essentiality score, while in contrast, the shRNA screen identified *WRN* as an essential gene with a negative essentiality score. We found multiple studies where non-pooled functional shRNA or drug screens were conducted, consistent with the results of shRNA screening but not CRISPR (Aggarwal *et al.*, 2013; Fausti *et al.*, 2013). As *WRN* is known to play critical roles in DNA repair, the essentiality of a *WRN* mutation is therefore likely to be interfered with the NEJM pathways triggered by CRISPR, which may explain the missing essentiality in the CRISPR screen.

**Fig. 6.**
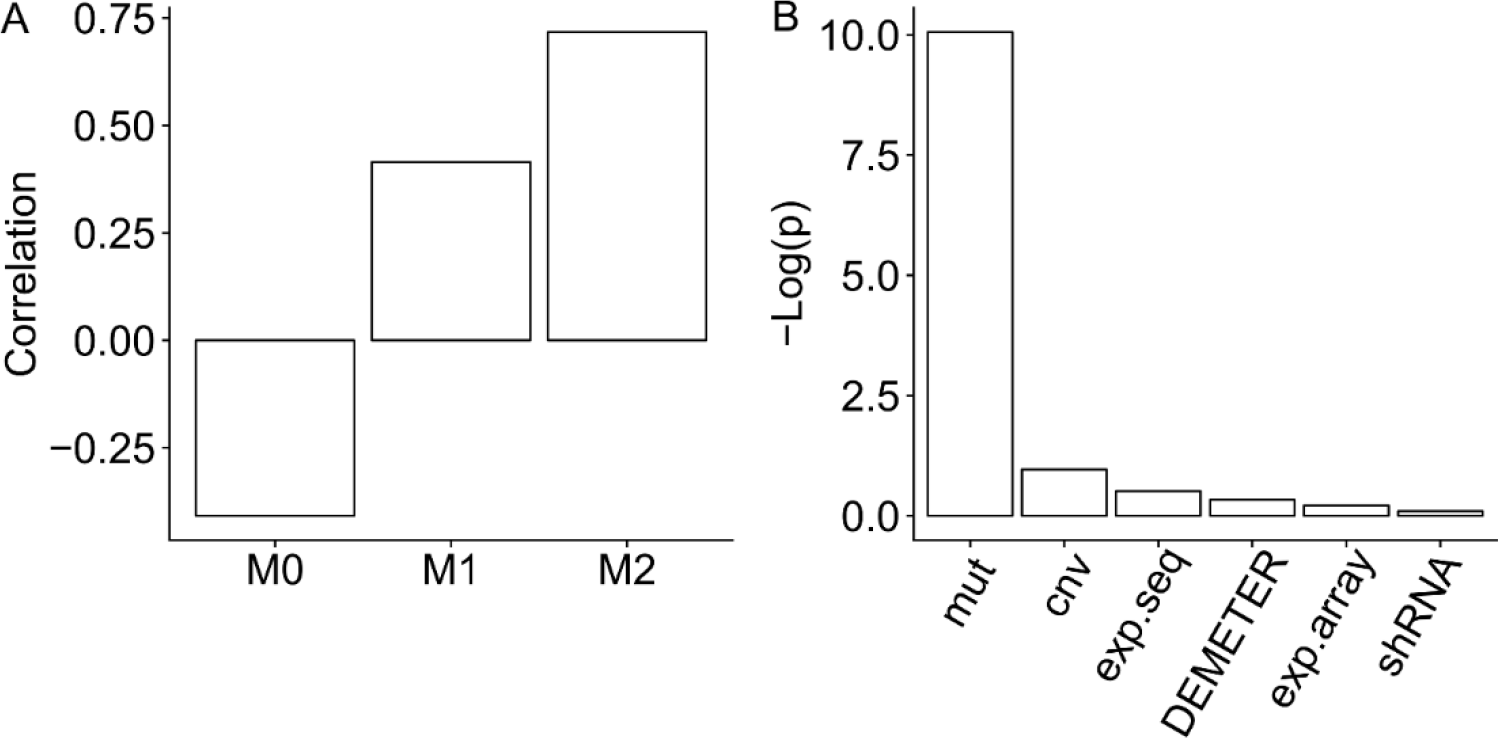
Model prediction performance for the *WRN* gene. (A) The correlations between the CRISPR essentiality score and the shRNA essentiality score (M0), and the predicted essentiality scores by M1 and M2, respectively. (B) The significances of the regression coefficients for the individual features in M2.

We summarized the counts of genes where the regression coefficients were significant using nominal p-value < 0.05 as the threshold. Consistent with our previous results, there were only 6.2% and 6.8% of the total genes where shRNA-based essentiality scores (shRNA and DEMETER, separately) were identified as significant features to predict the CRISPR score. Interestingly, we found that 13.4% of cases (n = 1,776) showed significant coefficients for copy number variation, which were much higher than the frequencies for the other feature types (mutation: 6.8%, sequencing-based gene expression: 8.0%, microarray-based gene expression: 7.3%). Of all the n = 1,776 genes with significant coefficients for copy number variation, we found that 76% of them (n = 1,420) have negative coefficients, indicating that an increase of copy number tends to reduce the CRISPR but not the shRNA essentiality score. This observation suggested that the copy numbers of the genes strongly influence the essentiality profile in the CRISPR screen, while such phenotypes could not be replicated in the shRNA screen. This observation was consistent with recent studies which showed that a CRISPR targeting genomic regions of high copy numbers may trigger unwanted DNA-damage responses, resulting in false positive of gene essentiality (Meyers *et al.*, 2017). Since cancer cell lines typically contain more genomic amplifications than normal cells, the CRISPR-based essentiality scores for genes in the amplified region should be interpreted with caution.

### Combined gene essentiality scoring helped identify core essential as well as cell-specific essential genes

Supported by the importance of molecular features for understanding the consistency between shRNA and CRISPR screens, we defined a combined essentiality score (CES) for a gene i in cell line j. We hypothesized that the CES score should resemble the true gene essentiality score via a linear combination of these observed functional and molecular features, based on the M2 model as below.

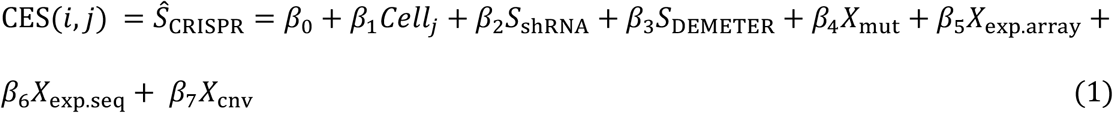

where β parameters are the regression coefficients that are fitted by M2 model using the least square estimates.

For each of the four versions of gene essentiality scores (i.e. CES, CRISPR, and the two variants of shRNA-bases scores denoted as DEMETER and shRNA), the ranking of genes in a given cell line can be determined. The rankings of a given gene across all the cell lines can then be averaged to represent the overall essentiality ranking of the gene. We defined the core essential genes as those with the overall ranking lower than 1,000 out of the 14,138 genes.

We compared the sets of core essential genes determined individually by these four scores. As shown in **Figure 7A**, the set of core essential genes based on the CES score (n = 252) has a limited overlap with those determined by CRISPR and shRNA scores. The minimal overlap was also observed between the CRISPR and shRNA-based core essential genes, corroborating the poor consistency that was found in the previous analyses. The majority of the CES-based core essential genes (n = 181 out of 252) was not detected by either CRISPR or shRNA-based screens, suggesting that the CES scores have identified novel essential genes after the inclusion of the molecular features.

**Fig. 7.**
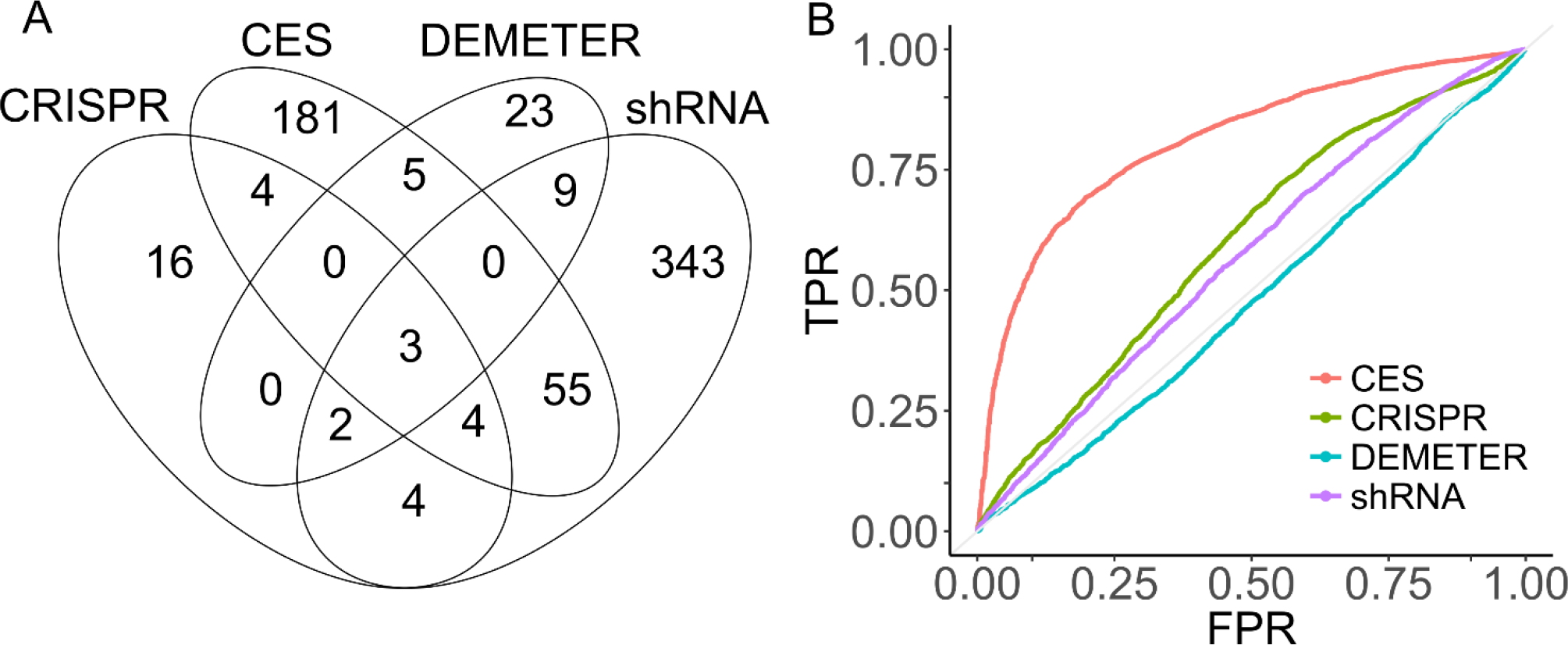
Performance comparison of different scoring method. (A) Overlap of the core essential genes identified by CRISPR, shRNA, DEMETER and CES. (B) The ROC curves for the prediction of housekeeping genes using the average ranking of the genes based on the four scoring methods, separately.

We investigated whether the average rankings based on these four scores can predict housekeeping genes, e.g. whether the housekeeping genes tend to be ranked higher in the list. By using the predefined set of housekeeping genes as the ground truth (n = 3,804, see Materials and Methods), we found that the CES score greatly outperformed CRISPR- and shRNA-based essentiality scores in term of ranking accuracy (AUROC = 0.81, as compared to 0.60 and 0.57 for CRISPR and shRNA scoring respectively, **Figure 7B**). Interestingly, The AUROC for the DEMETER-based ranking was lower than 0.5, suggesting that the DEMETER-corrected shRNA scores could not detect the majority of housekeeping genes consistently across the cell lines. Note that the DEMETER tends to attribute part of the essentiality to the off-target effects, and therefore may increase the false negative rate when detecting housekeeping genes.

We found that commonly known housekeeping genes were identified as core essential genes more often by CES than by CRISPR and shRNA scores. For example, *GAPDH* encodes the enzyme for regulating cell energy metabolism, the inhibition of which leads to cell apoptosis. *GAPDH* has been constitutively expressed at a high level in all cells and therefore is commonly used as the loading control for protein and gene expression experiments. In our study, *GAPDH* was ranked at the top 1000 for 38 cell lines using the CES scoring, while using the other scoring methods the gene was ranked at the top 1000 in less than 20 cell lines. The similar pattern was found for *ACTB* which is another highly conserved housekeeping gene (**Table 1**). These results suggested that CES is able to capture the true essential genes better than using CRISPR or shRNA-base screening data alone.

Finally, we utilized the CES score to identify cell-specific essential genes, which were defined as those ranked at the top 100 for a given cancer cell line, while their overall rankings across all the cell lines must go beyond the top 2000. To focus on those genes discovered by the CES scoring only, we removed those genes ranked at the top 5,000 by either CRISPR or shRNA-based scoring.

We found altogether 44 novel essential genes for 22 cell lines, for which the cell-gene mapping was shown in **Figure 8**. The majority of the genes were linked to only one cell line, as requested by the cell-specific selection criteria. For example, *AGR2* has been identified as an essential gene for the T47D cell line (breast cancer). *AGR2* has been reported to play critical roles in estrogen receptor (ER) positive breast cancer development. However, both the CRISPR and shRNA screens failed to identify *AGR2* as an essential gene for T47D cell line (essentiality score = 0.24 and 0.35, respectively). On the other hand, certain genes were shared by multiple cell lines, suggesting an extended level of specificity, which might allow them to be used as predictive biomarkers for disease stratification. For example, lysozyme encoded by the *LYZ* gene has been linked to four leukemia cell lines (MV411, MOLM13, OCIAML2 and MONOMAC1). Interestingly, these cell lines all belong to the AML (acute myeloid leukemia) subtype, suggesting the biological relevance of using *LYZ* as a drug target for AML. The actual functions of *LYZ*, together with other novel essential genes that were found specifically for certain groups of cell lines, might be worth further investigation to facilitate drug discovery in personalized medicine.

**Table 1.** Count of cell lines for which *GAPDH* and *ACTB* were identified as essential.

|  | shRNA | DEMETER | CRISPR | CES |
| --- | --- | --- | --- | --- |
| *GAPDH* | 18 | 14 | 17 | 38 |
| *ACTB* | 23 | 8 | 13 | 40 |

**Fig. 8.**
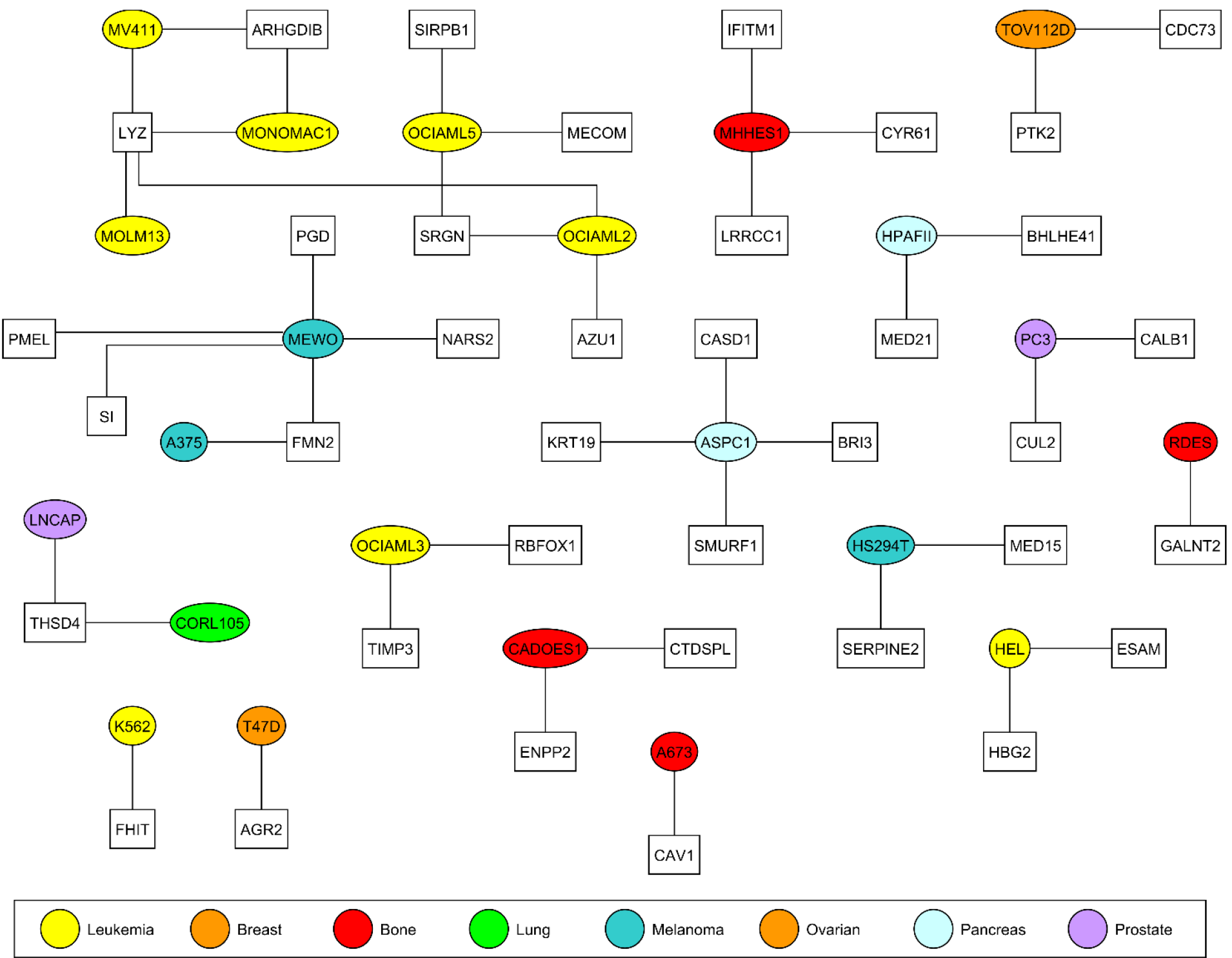
The novel cell-specific essential genes detected by CES. Round nodes are cell lines and square nodes are genes.

## Discussion

Loss-of-function screens using shRNA and CRISPR-based techniques have been commonly utilized for studying cancer dependency at the genome-scale, while questions remain on whether they produce consistent gene essentiality profiles for a given cancer sample. A recent side-by-side comparison of the K562 cell line (leukemia) observed a lack of consistency for the essential genes (Morgens, et al., 2016). In this study, we did a more systematic comparison using a panel of 42 cancer cell lines representing 10 tissue types, and confirmed that the inconsistency can be observed more generally across different cellular contexts. Furthermore, we found that the off-target effects in the shRNA screens could not explain the inconsistency, suggesting the inherent biases of these two technologies, which might be introduced by multiple confounding factors that are related to molecular features of the genes. Following this line, we proposed a multiple linear regression model called M2 to incorporate the molecular profiles of the genes. We showed that including molecular features as covariates in the M2 model did improve the consistency between shRNA and CRISPR-based essentiality scores. Interestingly, we found that copy number variation was among the top molecular features that may explain the difference between the two screen techniques. Indeed, recent studies showed that CRISPR screen may erroneously identify genes in copy-number-amplified region as essential, as DNA damage response and cell cycle arrest will be triggered independent of targeting genes (Fellmann *et al.*, 2017), while shRNA screen could not detect essential genes expressed at low levels (Hart *et al.*, 2015). These evidence prompted us to build a data harmonization model to incorporate molecular features to define a Combined Essentiality Score (CES). We showed that the CES harmonization greatly improved the consistency of gene essentiality profiles and thus helped to identify more essential genes for cancer cells.

Our pan-cancer essentiality result should provide a more convincing basis for subsequent drug discovery studies. The linear regression model is easy to apply and interpret. Other prediction methods, such as Support Vector Machine and Random Forest, however, will also be explored to further improve the prediction accuracy. Our method provides a novel perspective to explore the larger feature space, allowing an improved prediction of essential genes at individual cell lines. The effective integration of functional and molecular data might provide important clues for the development of targeted drugs directed towards personalized medicine.

Our method is not limited to interrogating essential genes, but also applicable for discovering genes that are correlated with other complex cell phenotypes. However, due to the scarcity of such data, we will keep this direction as a future step. We have not evaluated the other variants of CRISPR-Cas9 based screens such as CRISPRi and CRISPRa. On the other hand, the siRNA (small interference RNA) technique has been considered more suitable for array-based assays in a non-pooled fashion (Mullenders and Bernards, 2009), and therefore was not considered in the current study even though it allows more complex phenotypes to be measured. The pooled functional screens are largely restricted for simple phenotypes such as cell viability and toxicity. In comparison, arrayed screens allow for complex molecular readouts such as transcriptional responses, but usually with only one gene depletion at a time. Recent technology development in single cell sequencing has enabled the testing of multiple phenotypes for specific gene depletions in the pooled screens (e.g. CROP-seq and Perturb-seq) and we expect that our method can be also applicable there (Datlinger *et al.*, 2017). Systematic comparisons among the other variants of functional screen techniques should be investigated further.

In addition to human cancer cell lines that were studied here, poor phenotypic consistencies were also observed in other model organisms including mouse and zebrafish (Kok *et al.*, 2015; Young *et al.*, 2009). We foresee that our data harmonization model and the CES scoring might be extended to improve the consistency of shRNA and CRISPR screens in non-cancer cell lines and model organisms.

## Acknowledgements

We thank the researchers, especially those from Broad Institute (project Achilles and project CCLE), for making their data publicly available. This work was supported by funds from European Research Council Starting Grant agreement No 716063 (DrugComb) and Helsinki Institute of Life Sciences Research Fellow funding. W.W and A.M. are supported by the FIMM-EMBL PhD program.

*Conflict of Interest:* none declared.

